# Intrinsically disordered protein’s coil-to-globule transition and adsorption onto a hydrophobic surface under different conditions

**DOI:** 10.1101/2023.03.08.531675

**Authors:** Bernat Durà Faulí, Valentino Bianco, Giancarlo Franzese

## Abstract

Intrinsically disordered proteins (IDPs) and proteins with intrinsically disordered regions (IDRs) can modulate cellular responses to environmental conditions by undergoing coil-to-globule transitions and phase separation. However, the molecular mechanisms of these phenomena still need to be fully understood. Here, we use Monte Carlo calculations of a model incorporating water’s effects on the system’s free energy to investigate how an IDP responds to a hydrophobic surface under different conditions. We show that a slit pore confinement without top-down symmetry enhances the unfolding and adsorption of the IDP in both random coil and globular states. Moreover, we demonstrate that the hydration water modulates this behavior depending on the thermodynamic parameters. Our findings provide insights into how IDPs and IDRs can sense and adjust to external stimuli such as nanointerfaces or stresses.

## Introduction

Structured proteins fold into a specific three-dimensional structure to achieve their function. However, proteins with intrinsically disordered regions (IDRs) and intrinsically disordered proteins (IDPs) have regions or domains that remain unfolded or disordered under physiological conditions. IDRs larger than 30 amino acids and IDPs are common in cells and regulate diverse cellular processes, such as RNA binding, oligomerization, metabolite recruitment, and catalysis.^1^ Moreover, IDRs and IDPs are exposed to weak, multivalent, and dynamic interactions that could lead to liquid-liquid phase separation (LLPS), a phenomenon in which they form droplet-like structures that concentrate biomolecules without a membrane barrier. The biomolecular condensation potentially involves various biological functions and dysfunctions, such as gene regulation, signal transduction, and neurodegeneration. ^2,3^ Interestingly, IDRs and IDPs can phase-separate at much lower concentrations than structured proteins, such as those involved in cataract formation or fibrils. However, the balance between liquid-like and solid-like phases is delicate and depends on the type of interaction among the disordered molecules. For example, homotypic interactions tend to promote aggregation and fibrillation, which can be detrimental to cellular health. On the other hand, heterotypic interactions can stabilize the liquid phase and prevent pathological phase transitions. ^4^

Recent studies have linked IDPs’ coil-to-globule transition to their LLPS. This allows for sequence-specific phase diagrams to be calculated. ^5^ It is, therefore, interesting to explore how heterotypic interactions of IDPs can affect their coil-to-globule transition (and condensation) using simple models.

Furthermore, in many fields like medicine,^6–9^ food science,^10–12^ and biosensors,^13–15^ it is essential to understand how proteins and biomolecules interact with nanomaterials. For example, when nanoparticles come into contact with bloodstreams, they form a corona of multiple layers of proteins and biomolecules. This gives the nanocomplex a new biological identity.^16,17^ It is generally accepted that, upon adsorption, proteins can alter their structure,^18–20^ which can have significant consequences like an inflammatory response or fibril formation.^21,22^ However, our comprehension of these mechanisms must still be completed.^23^ Also, the effect of adsorption on a flat surface can be highly diverse when comparing structured regions with IDRs of the same protein. ^24^ Hence, understanding the impact of the interface on the protein’s conformation is crucial in determining nanomaterial interactions with biological environments.^17,25–29^

Here we consider the coarse-grained Bianco-Franzese (BF) model for proteins in explicit water in its simplest version,^26,30^ as defined below. Despite its schematic approximations, the BF model can show, both for structured proteins with a native state and for IDPs, that accounting for the contribution of the hydration water^31^ is enough to predict protein thermodynamic properties consistent with theories^32,33^ and experiments.^34,35^

The BF Hamiltonian model reproduces elliptically shaped protein stability regions (SR) in the temperature-pressure (*T* -*P*) plane.^36,37^ It includes high-*T* unfolding (melting), driven by the entropy increase, which is common to all the protein models, e.g., Ref. ^38^ Additionally, it shows that hydration water energy drives the low-*T* (cold) unfolding. Hydrophobic-collapse models cannot explain this experimental phenomenon. ^39,40^ Specific models can reproduce the cold unfolding without^41,42^ or with^43,44^ a *P* -dependent behavior. However, at variance with the BF model, they do not reproduce the experimental elliptic SR.

Moreover, the BF model explains high-*P* unfolding as density-driven due to increased hydration water compressibility at hydrophobic interfaces, ^45–48^ common also to other water-like models.^49^ Finally, it explicates the low-*P* denaturation seen in the experiments^37,50^ and models^51^ as enthalpy-driven.^30^

The BF model has other interesting properties. For example, it sheds light on water’s evolutionary action in selecting protein sequences and the effect of extreme thermodynamic conditions. This has implications for protein and drug design.^52^ For example, the model shows that artificial covalent bridges between amino acids are necessary to avoid protein denaturation at *P >* 0.6 GPa.^37^ Moreover, it also helps us understand why only about 70% of the surface of mesophilic proteins is hydrophilic, and about 50% of their core is hydrophobic.^52^

Recently, the BF model has been used to study how structured proteins denature and aggregate reversibly depending on their concentration in water solutions with one^53^ or two protein components^54^ or near hydrophobic interfaces. ^27^ The results show that unfolding facilitates reversible aggregation^53^ with a cross-dependence in multicomponent mixtures.^54^ Also, the proteins aggregate less near hydrophobic interfaces, at high *T*, or by increasing the hydrophobic effect (e.g., by reducing salt concentration). ^27^

Here, we study by Monte Carlo calculations how adsorption on a hydrophobic interface affects the coil-to-globule transition of an IDP. The results help us to understand the fate of proteins once they interact with nanomaterials^55,56^ and the possible effects for proteins undergoing phase separation.

## Model

### The FS model for water

The BF model is based on adding a coarse-grained protein with its hydrated interface to the Franzese-Stanley (FS) water model. ^57–60^ The FS model includes cooperative (many-body) interactions in an elegant model proposed by Satsry et al.^61^ with only two free parameters. The FS model adds a third parameter describing the hydrogen bonds (HBs) cooperativity.

The new parameter quantifies the relative strength of many-body HBs compared to van der Waals interactions and controls the phase diagram in the supercooled region. ^62^

The FS model coarse grains the water atomistic coordinates, introducing a density field with local fluctuations due to the HB structure but keeping a molecular description of the HB network. Recent reviews summarize the definition of the FS model for a water monolayer and its main properties.^63,64^

The extension of the FS model to bulk shows that its three parameters can be adjusted in a way to give optimal agreement with the experimental water data in an extensive range of *T* and *P* around ambient conditions^65^ (for preliminary calculations, see Ref.^66^). However, the HB network’s peculiar structure that preferentially has a low (four) coordination number makes the monolayer version of the model, with only four neighbors, interesting. Indeed, the FS monolayer offers a reasonable coarse-grained approximation for water near ambient conditions at the cost of renormalizing its parameters. This renormalization allows us to account for the difference in entropy compared to the bulk, with the advantage of being easier to visualize and calculate.

Therefore, we consider a partition of the system’s two-dimensional projection into *N* square cells, of which water molecules occupy *N*_*W*_ ≤ *N*, each with the average volume *v*(*T, P*) ≥ *v*_0_, the van der Waals excluded volume for a water molecule without HBs. On the other hand, we assume that the HBs are the primary source of local density fluctuations and associate with each HB a proper volume *v*_HB_*/v*_0_ = 0.5 equal to the average volume increase per HB between high-density ices VI and VIII and low-density (tetrahedral) ice Ih, approximating the average volume variation per HB when a tetrahedral HB network is formed.^67^ Hence, the volume of water is

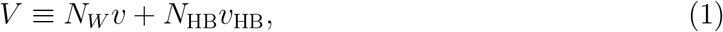

where *N*_HB_ is the number of HBs.

The FS Hamiltonian, describing the interaction between the water molecules, is

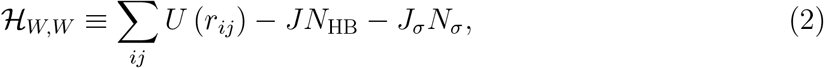

where *U* = ∞ for 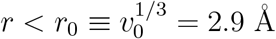, and 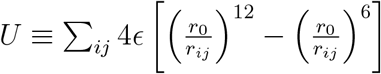 for *r*_0_ *< r <* 6*r*_0_ (cutoff) or *U* = 0 for larger *r*, with *ϵ* = 5.8 kJ/mol. The sum runs over all possible water-molecule couples (including those in the hydration shell introduced in the BF model). This term accounts for the O–O van der Waals interaction between molecules *i* and *j*. Previous calculations prove that the model results are independent of the details of *U*.^68^

Because we consider the system at constant *NPT*, the distance *r*_*ij*_ is a continuous variable. Notably, because the formation of HBs does not change the nearest neighbor (NN) distance 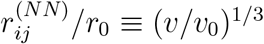 between water molecules in the first coordination shell,^67^ the van der Waals interaction is unaffected by the HBs, guaranteeing that the FS is not just a simplified mean-field model.

The term −*JN*_HB_ accounts for the additive (two-body) component of the HB. The FS model adopts the HB definition based on the distance between the centers of mass of two water molecules and the angle between the OH group of one and the O atom of the other^57^ The HB has minimum energy when the H is along the O-O direction or deviates less than 30^*o*^.^64,69,70^ Hence, only 1/6 of all the possible orientations in the plane of the H atom relative to the O-O direction corresponds to a bonded state, while the other 5/6 states are nonbonded. Therefore, to correctly account for the entropy variation once the HB is formed, we introduce a 6-state bonding variable *σ*_*ij*_ for each of the four possible HBs that each water molecule *i* can form with a NN water molecule *j*. We assume that the HB is formed only if both molecules have the same bonding state, i.e., if 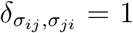, where *δ*_*ab*_ = 1 if *a* = *b*, 0 otherwise.

On the other hand, the HB can be considered broken when the O-O is larger than a given *r*_max_.^71^ The FS model assumes the reasonable value *r*_max_ ≃ 3.65Å,^64^ implying that for *r > r*_max_ it is (*r*_0_*/r*)^3^ ≡ *v*_0_*/v <* 0.5. Hence, by setting *n*_*i*_ = *n* ≡ *θ*(*v*_0_*/v* − 0.5), where *θ*(*x*) is the Heaviside step function, the total number of HBs is 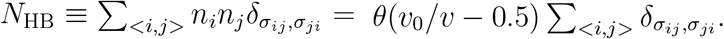.

The last term in the Hamiltonian −*J*_*σ*_*N*_*σ*_ accounts for the many-body term that can be calculated by *ab initio* methods. It favors the formation of a low-density (tetrahedral) local structure in liquid water even at ambient conditions.^72^ In classical atomistic potentials, this term is modeled with a long-range polarizable dipolar interaction. However, recent calculations, based on polarizable models including the MB-pol potential,^73–76^ show that it can be approximated with a short-range 5-body interaction within the first coordination shell of a water molecule.^77^ This result gives a solid theoretical foundation to the FS assumption of modeling the cooperative term as an effective 5-body interaction within the first coordination shell of each water molecule *i*, with 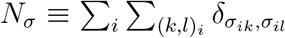, where the inner sum is over all the pairs of the bonding variables of the molecule *i*. Following,^30^ we set here *J/*4*ϵ* = 0.3 and *J*_*σ*_*/*4*ϵ* = 0.05.

### The BF model for a hydrophobic IDP

Based on atomistic results, the BF model assumes that in a hydrophobic (*ϕ*) hydration layer (Fig. 1)

**Figure 1:**
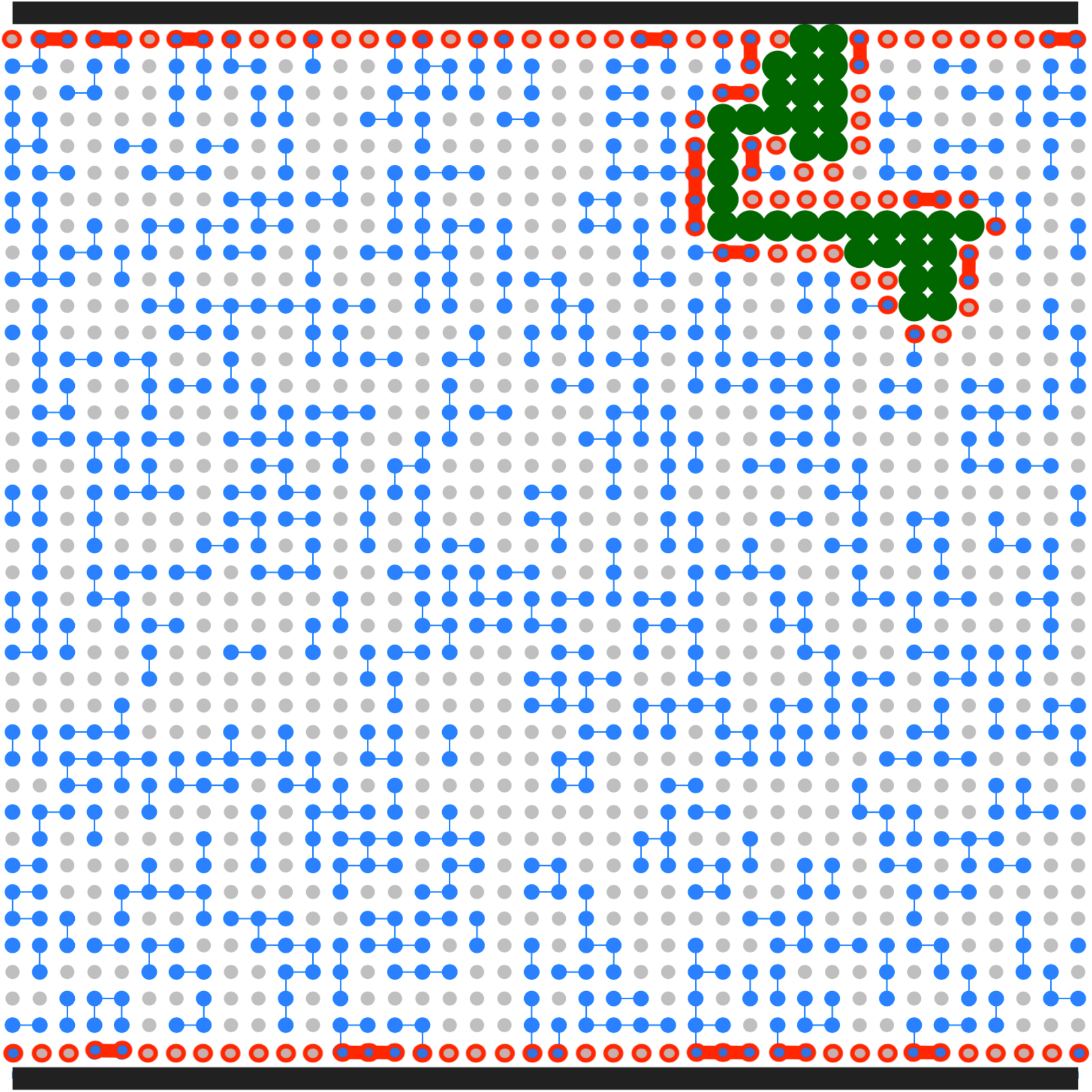
Example of one of the visited conformations for an IDP confined in a slit-pore. The IDP, coarse-grained in the BF model as a chain of (green) beads, is adsorbed onto one of the pore’s walls (black top and bottom lines) and surrounded by water. At the thermodynamic conditions (*k*_*B*_*T/*4*ϵ*=0.55 and *Pv*_0_*/*4*ϵ*=0.4) of this example, some water molecules (grey dots) do not form any HB, while others (blue dots) can have up to four HBs (blue lines). Water molecules and water-water HBs in the (protein and wall) hydration shells are highlighted in red.

i. the interfacial water-water HBs are stronger than bulk HBs, with an extra interaction Δ*J*^(*ϕ*)^*/J* = 0.83;
ii. the water compressibility is larger than bulk compressibility so that HB’s volume is reduced by 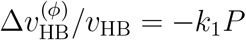, with *k*_1_ = *v*_0_*/*4*ϵ*.

Hence, the FS enthalpy 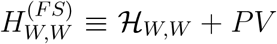, from Eq.s (1, 2), acquires an extra term in the BF model that for the hydrophobic hydration shell is

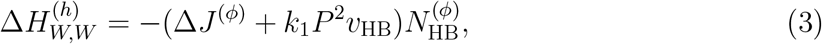

where 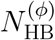 is the number of HBs between water molecules in the hydration shell.

For the proteins, the BF model adopts a coarse-grain representation of beads-on-a-chain, with one bead per residue, that has been extensively used in the literature to get a qualitative understanding of protein properties. ^52^ The protein Hamiltonian

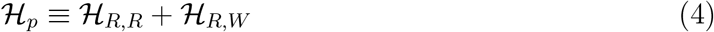

describes the interactions among the protein NN residues, *ℋ*_*R,R*_, and between the residues and the NN water molecules in the hydration shell, *ℋ*_*R,W*_.^52^ Here we represent the IDP with a hydrophobic homopolymer where all the *N*_*R*_ residues interact with the NN molecules by excluded volume. A more general expression for *ℋ*_*p*_ accounting for the complete protein amino acids is presented in Ref.s.^30,35,52^

Finally, the BF enthalpy of the entire system with the hydrated protein in explicit water is

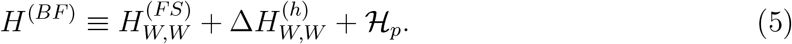

The general expression for the Gibbs free energy of the BF model is

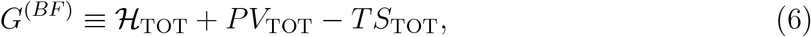

where 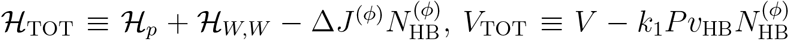, and *S*_TOT_ is the total entropy of the system associated with all the configurations with the same number of proteins contact points (CPs), *N*_CP_, defined in the following, and the same number of water molecules in the hydration shell (red dots in Fig. 1).

As in the BF original formulation, we assume that protein residues and water molecules have the same size. This simplification could be removed by letting each residue occupy several cells, eventually increasing the hydrated protein surface. However, we expect that, by rescaling the hydration parameters, there would be no qualitative change in our findings.

### Monte Carlo calculations with and without top-down symmetry

We realize the slit pore geometry in a square partition with size *L* = 40 by fixing *L* hydrophobic cells along a line and applying periodic boundary conditions in all directions (Fig. 1). We perform Monte Carlo (MC) calculations for a protein with *N*_*R*_ = 36 residues at constant *P, T, N*_*W*_ and *N*_*R*_, with *N*_*W*_ + *N*_*R*_ = *N* ≡ *L*^2^.

We consider random initial configurations and equilibrate the water bonding indexes with a clustering algorithm^78^ and the protein chain with corner flips, pivots, crankshaft moves, and random unitary translations of its center of mass. ^79^ A single MC step is made of a random sequence of move attempts for each degree of freedom of the system (36 residues and 6256 *σ*_*ij*_ variables). After moving a protein, the cells left by the amino acids are replaced by water molecules whose values of the four *σ*_*ij*_ variables are chosen randomly.^27,33,80^

To facilitate protein adsorption, we break the top-down symmetry by biasing the translation toward one of the confining walls but not along the slit pore. For the sake of the description, we call *top* the biased wall. The bias mimics a drift or a weak force pushing the protein toward the top interface without limiting its thermal motion parallel to the walls. In the Supplementary Material, we discuss the case without bias, i.e., with top-down symmetry. We perform calculations for temperatures ranging from *k*_*B*_*T/*4*ϵ*=0.01 to 0.6 and pressures from *Pv*_0_*/*4*ϵ* = −0.2 to 0.6. For each (*T, P*), we collect configurations for every 100 of 10^6^ MC steps after discarding 10^4^ equilibration steps.

Our main observable is how close the IDP is to a globule conformation. To this goal, we calculate the degree of folding of the protein as the number *N*_CP_ of contact points (CP) that the protein has with itself. We consider that there is a CP if two residues of the protein occupy the NN cells but are not adjacent along the chain.

## Results and discusion

### Coil-to-globule transition

For each (*T, P*), we compute the average *N*_CP_ for the IDP. For our 36 residue-long IDP, the maximum *N*_CP_ is 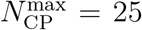. When 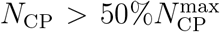, we identify the IDP conformation as globule, while we consider it coil otherwise (Fig.2). Our calculations show that, for *Pv*_0_*/*4*ϵ <* 0.5, *N*_CP_ is non-monotonic as a function of *T*. For *Pv*_0_*/*4*ϵ* = 0.3 (blue line in Fig.2), *N*_CP_ is larger than 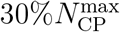 in a limited range of *T*, but does not reach the 40% threshold. Within our resolution of *P, Pv*_0_*/*4*ϵ* = 0.2 (yellow line in Fig.2) is the highest at which the IDP reaches 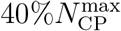, while for any *Pv*_0_*/*4*ϵ* ≤ 0.1 (orange line in Fig.2) the IDP undergoes a coil-to-globule transition.

**Figure 2:**
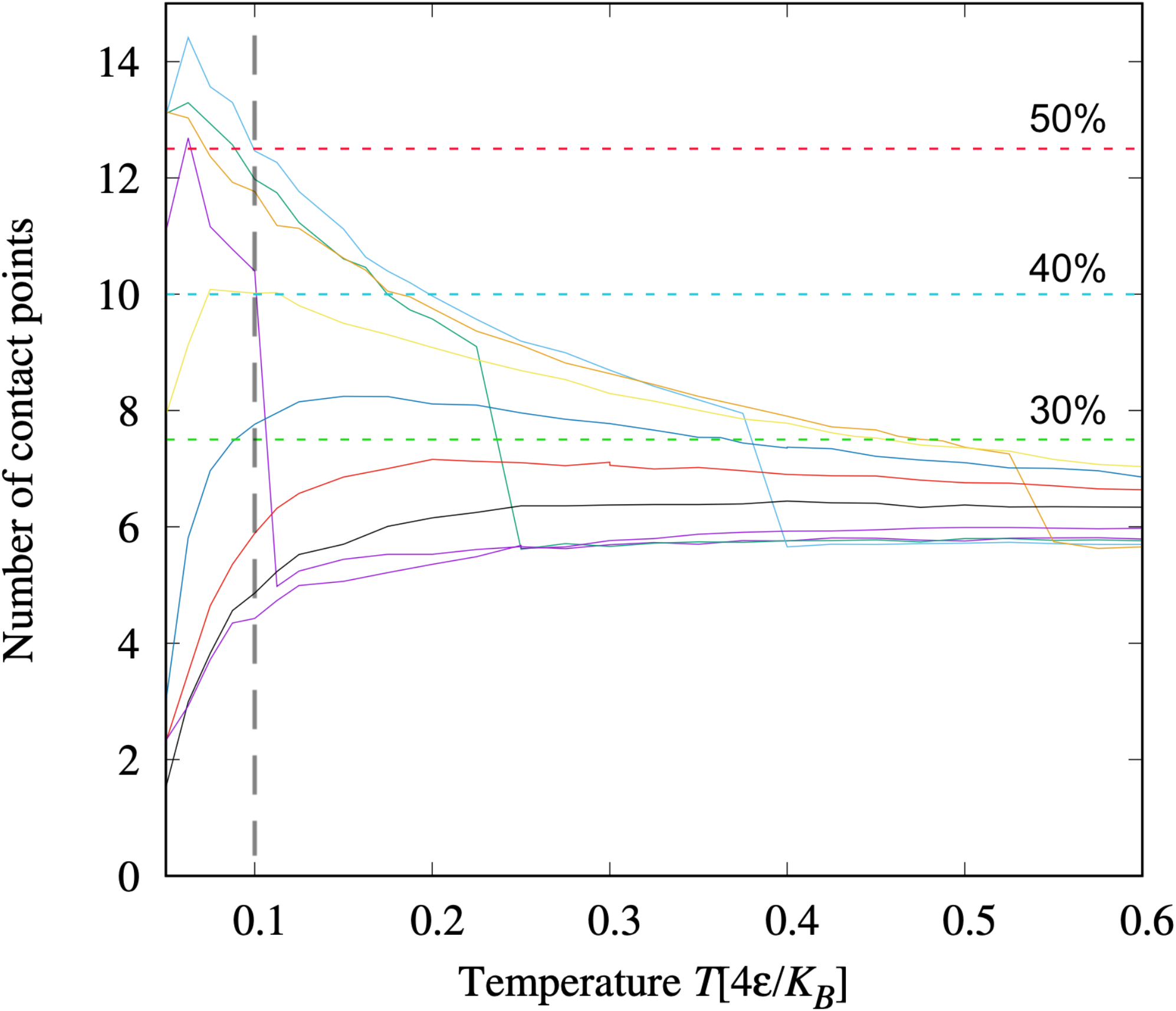
Coil-to-globule transition for the IDP in the hydrophobic slit pore with-out top-down symmetry. The number of CPs, *N*_CP_, at constant *P* as a function of *T* is non-monotonic for any *Pv*_0_*/*4*ϵ <* 0.5, showing a reentrant coil-to-globule transition when we consider CP’s thresholds at 50%, 40%, and 30% of CP_max_ (red, blue, and green horizontal dashed lines, respectively). The calculations are presented as segmented lines (points connected by segments) for pressures, from bottom to top at *k*_*B*_*T/*4*ϵ* = 0.1 (vertical dashed grey line), *Pv*_0_*/*4*ϵ* = 0.6 (indigo), 0.5 (black), 0.4 (red), 0.3 (blue), 0.2 (yellow), -0.2 (indigo), 0.1 (orange), -0.1 (green), 0.0 (turquoise). Note that the negative pressures intercalate with the positive. Discontinuities for the four lowest pressures mark the limit of the liquid-to-gas spinodal of the confined water solution when *T* increases.

These results are summarized in the *T* –*P* thermodynamic plane as stability regions (SRs) (Fig. 3). We find that the SRs at 30%, 40%, and 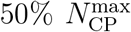 are concentric as expected.^30^ The three SRs display a reentrant behavior in *T* at different *P*, while the SR at 30% also shows a reentrant behavior in *P* at different *T*. Each SR line can be adjusted to curves with different degrees of ellipticity, as expected by general arguments.^32^ All the curves intersect the limiting temperature *k*_*B*_*T/*4*ϵ* ≲ 0.05 below which we cannot equilibrate the system within our statistics (the grey region in Fig. 3). Moreover, the SR for 30% intersects the liquid-to-gas spinodal line for our confined water solution. This line is marked by a significant volume increase of the entire system (not shown) and by discontinuities in *N*_CP_ for the four lowest pressures, reaching values typical of a random coil as at high *P* (Fig.2).

**Figure 3:**
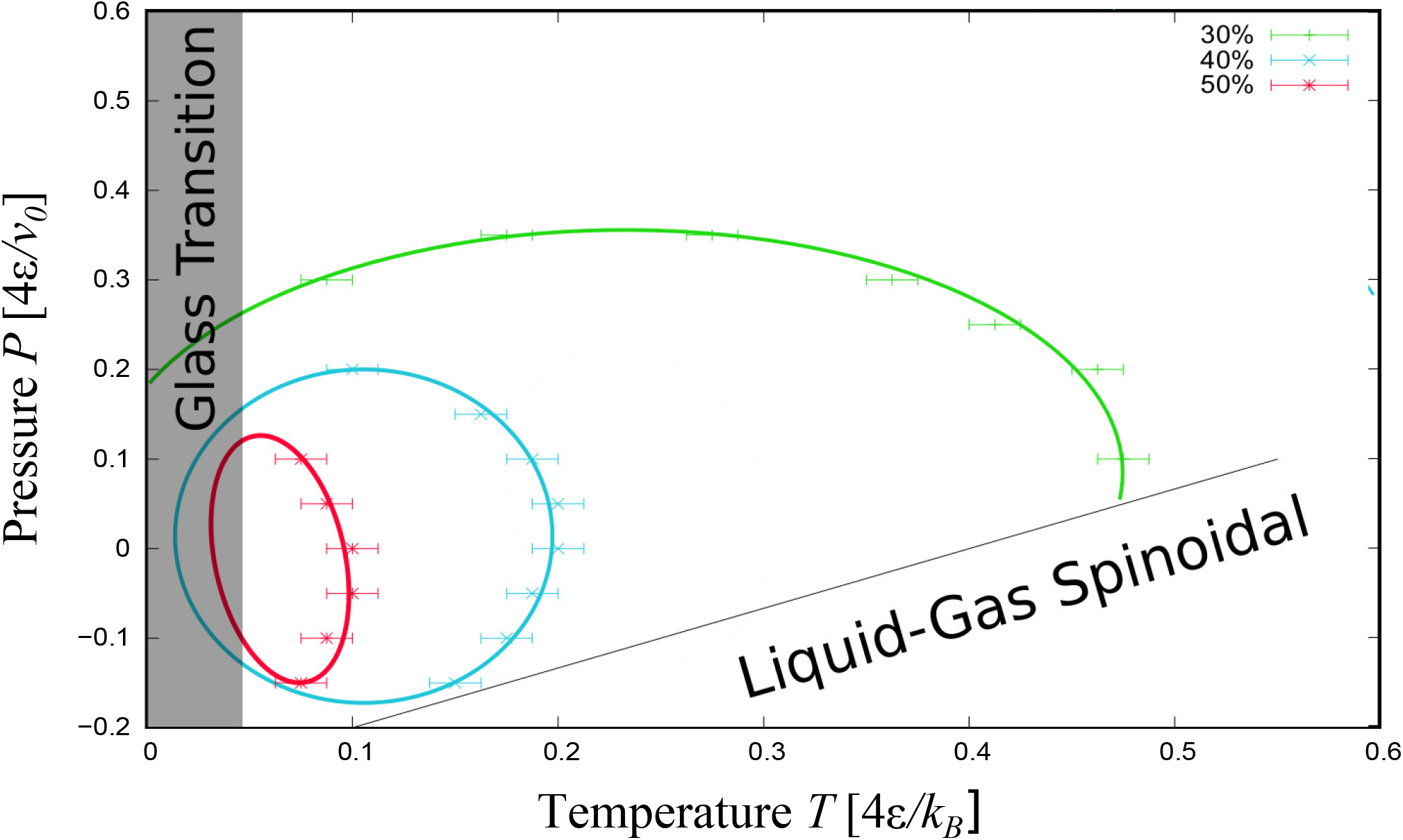
Stability regions for the IDP in the hydrophobic slit pore without topdown symmetry. Green, blue, and red symbols with error bars mark the state points where the IDP has, on average, *N*_CP_ *>* 30%, 40%, and 50% 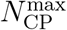, respectively. Elliptic lines are a guide for the eyes. The black line marks the liquid-to-gas spinodal for the confined water solution. The grey region indicates the glassy state points at *k*_*B*_*T/*4*ϵ* ≲ 0.05.

### Comparison with the transition without the slit pore

The hydrophobic confinement without top-down symmetry affects both the water and the protein. It changes the limit of stability (spinodal) of the liquid water compared to the gas phase (Fig. 4). At fixed pressure, we find the spinodal at lower *T* than the free IDP case.^30^ Overall the new spinodal is parallel to the former with a shift to lower *T* of ≃ 0.5*k*_*B*_*T/*4*ϵ* at constant *P*. This effect is independent of breaking the top-down symmetry (Fig. S1).

**Figure 4:**
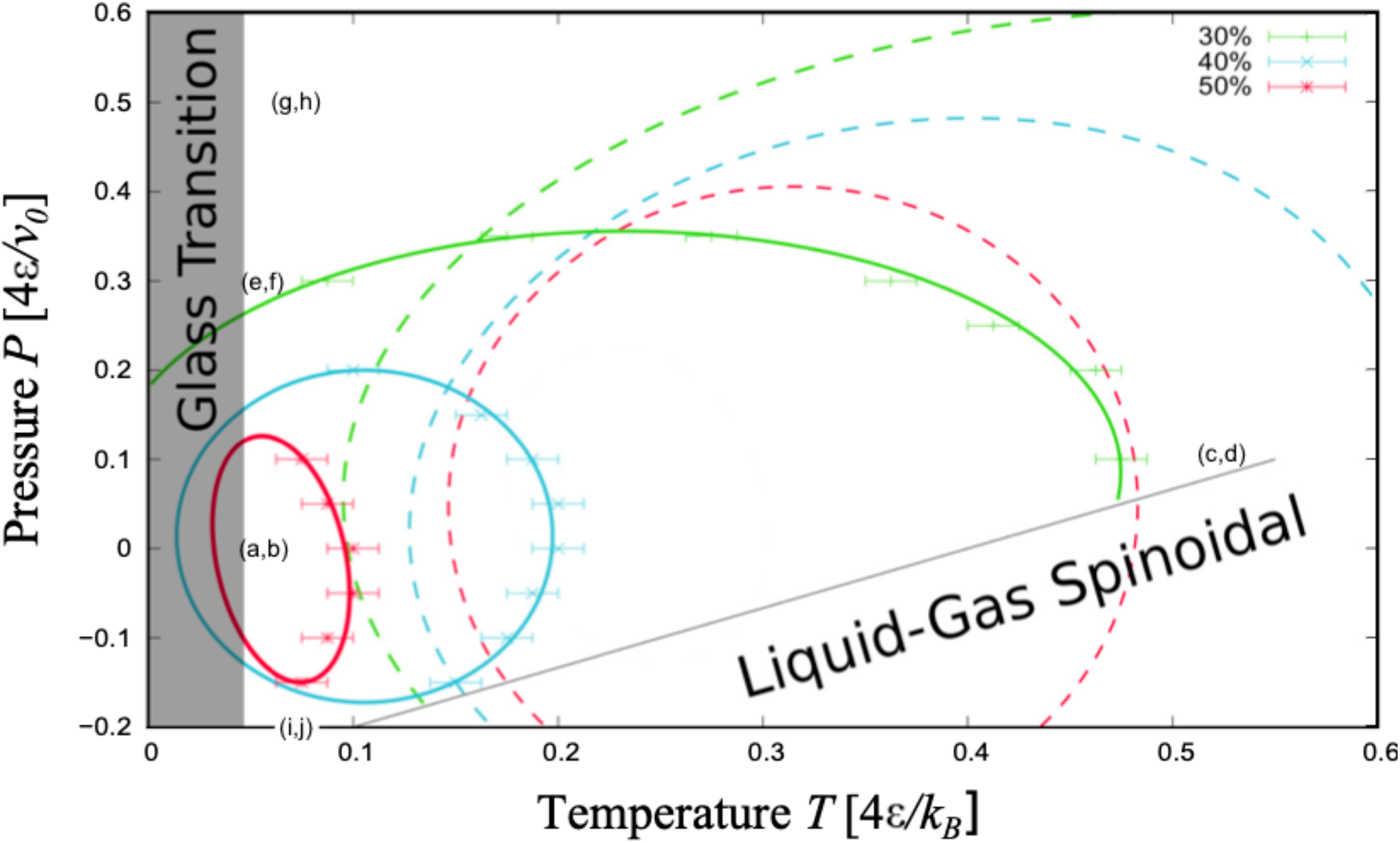
The hydrophobic confinement without top-down symmetry destabilizes the IDP globular conformations compared to the bulk case and affects the water liquid-to-gas spinodal. The latter (continuous black line) is shifted, at constant *P*, to lower *T* by ≃ 0.5*k*_*B*_*T/*4*ϵ* relative to the bulk case (not shown here because out of scale). The 30%, 40%, and 50% SRs for the IDP (continuous lines as in Fig 3) are displaced to lower (*T, P*) and are smaller compared to those for the free IDP (dashed lines with the same color code as the continuous). The grey area is as in Fig 3. The labels (a,b), (c,d), etc., refer to the state points discussed in Fig. 5. All the lines are guides for the eyes. The dashed lines are adapted with permission from Ref.^30^ copyrighted (2015) by the American Physical Society https://doi.org/10.1103/PhysRevLett.115.108101.

This result is a consequence of the interaction of the liquid with the slit pore. Confinement generally affects the properties of liquids, particularly water.^81–86^ The effect of hydrophobic evaporation has been extensively studied for confined water, e.g., in Ref.^87^ and references therein. Near ambient conditions, water dewets the walls of a hydrophobic nano-pore and evaporates.^88^ Experiments show capillary evaporation at scales consistent with the size of our slit pores at lower *P* and higher *T* compared to ambient conditions.^89^

Moreover, the presence of hydrophobic walls without top-down symmetry modifies the protein’s SRs compared to the free case (Fig.4). We find two striking features. First, all the regions marking 30%, 40% and 50% of 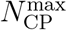 for the confined IDP occur at values of *T* and *P* that are lower than those for the free IDP.^30^ Second, the confined IDP has a coil-to-globule transition in a (*T, P*) range much smaller than the free case. ^30^ As a consequence of these changes, if a free IDP comes into contact with the biased hydrophobic surface at a thermodynamic condition where it is in a globule state, e.g., (*Tk*_*B*_*/*4*ϵ, Pv*_0_*/*4*ϵ*) = (0.4, 0.1), its *N*_CP_ would reduce from more than 50% to 30% of 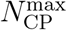 (Fig.4).

To check the effect of the bias, we repeat the calculations for a slit pore with top-down symmetry and find no differences for the confined water phase diagram, while we observe that the change in the protein stability compared to the bulk case is negligible (Fig. S1). Furthermore, to check the effect of the energy gain for the HB at the hydrophobic interface, we also decrease the Δ*J*^(*ϕ*)^*/J* parameters in the unbiased case (Table S1). This change implies that the protein SR shifts to lower *T* and is less accessible than the case with larger Δ*J*^(*ϕ*)^*/J*. Consequently, the unfolding at low *T* and low *P* are not observed for the weak Δ*J*^(*ϕ*)^*/J*.

Hence, our results suggest that facilitated adsorption, e.g., due to an attractive force or a drift toward the interface, destabilizes the globular conformations of the IDP significantly. As a further confirmation of this observation, we find that the maximum number of CPs that the protein reaches in the biased hydrophobic slit pore is 55% of 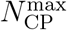, while it is more than 70% for a free IDP.^30^

### Interplay of adsorption and coil-to-globule transition

The thermodynamic state point affects not only the coil-to-globule transition but also the adsorption-desorption transition of the IDP on the hydrophobic surface, showing an intriguing interplay between the two phenomena.

#### Adsorption in the globule state

At low *T* and *P*, e.g., at (*k*_*B*_*T/*4*ϵ, Pv*_0_*/*4*ϵ*) = (0.050, 0.0), one expects that the most relevant term in the BF Gibbs free energy *G*^(*BF*)^, Eq.(6), is the interaction energy, *ℋ*_TOT_, while both *TS*_TOT_ and *PV*_TOT_ contributions are vanishing. Because *ℋ*_TOT_ is dominated by the *N*_HB_ term, the *G*^(*BF*)^ minimum corresponds to a maximum in *N*_HB_. Hence, the IDP adsorbs onto the surface, allowing more water molecules to form bulk HBs.

The water release induces, macroscopically, an effective hydrophobic attraction between the surface and the residues. As expected for the low relevance of the entropic and volumic terms in *G*^(*BF*)^ under these conditions, the many HBs organize in a highly ordered network with low *S*_TOT_ (Fig.5 a) and large volume (Fig.5 b), independent of the presence of bias (Fig. S2 a, b). In particular, for the unbiased case, the protein adsorbs onto the hydrophobic interface when it is in its globule state (Fig. S3).

This result is consistent with experiments. For example, blood proteins adsorb and form a corona onto nanoparticles with hydrophobic patches. ^90^ The common understanding is that the effect is maximum if the proteins flatten onto the nanomaterial. ^19,20^ Yet, experiments show that at least a part of the proteins in the corona can retain their functional motifs to allow the receptors’ recognition^91–93^ especially *in vivo*.^94^ In particular, the IDRs of structured proteins can be almost unaffected in their globular state when adsorbed onto a surface.^24^

Our results offer a rationale for this surprising experimental result. Indeed, we observe that the adsorbed IDP often keeps a globule conformation at *T* and *P* within its SR, as shown in movies mov1.mp4 and mov1nobias.mp4 in Supporting Information (SI) for biased and no-biased slit-pore, respectively. This is because *ℋ*_TOT_ is minimized when both *N*_HB_ and 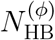, i.e., the number of HBs in bulk and within the hydration shell, respectively, are maximized. Hence, the IDP adsorps onto the surface to maximize *N*_HB_ but leaves as much as possible of the hydrophobic interface exposed to water to maximize 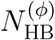 (Fig.s 5 a, and S2 a).

**Figure 5:**
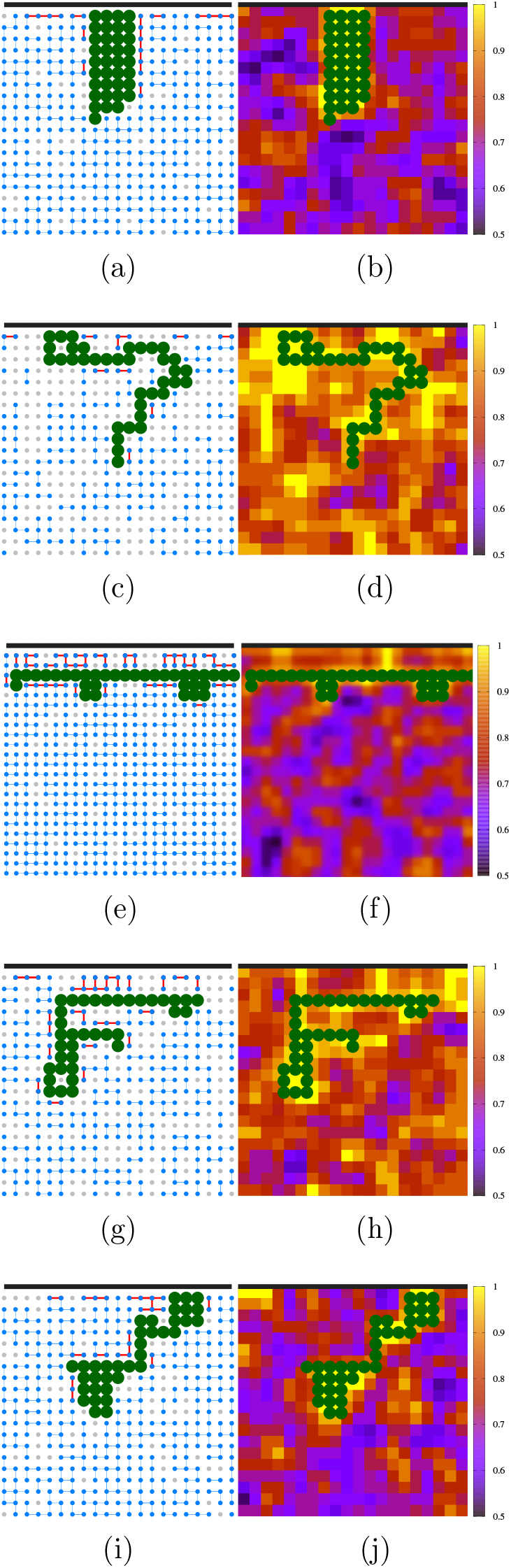
Coil-to-globule transition and adsorption without top-down symmetry at different state points. Left panels: HB network of bulk (blue) and hydration (red) water. Right panels: Color-coded water density field from low (dark blue) to high (yellow). The regions with no HBs (grey dots on the left panels) have a higher density (yellow regions on the right panels). The thermodynamic state points (*k*_*B*_*T/*4*ϵ, Pv*_0_*/*4*ϵ*) of the panels are reported in the phase diagram in Fig. 4: **(a), (b)** (0.050, 0.0); **(c), (d)** (0.525, 0.1); **(e)**,**(f)** (0.050, 0.3); **(g), (h)** (0.075, 0.5); **(i), (j)** (0.075, -0.2). The IDP and the interface are represented as in Fig. 1.

#### Adsorption in the coil state at high *T*

At higher *T*, approaching the liquid-gas spinodal, e.g., at (*k*_*B*_*T/*4*ϵ, Pv*_0_*/*4*ϵ*) = (0.525, 0.1), the entropy dominates the Gibbs free energy, Eq.(6), and the IDP loses its globule conformation, increasing *S*_TOT_ (Fig. 5 c, d for the biased case and Fig. S2 c, d for the unbiased). Most of the time, the IDP is kept adsorped onto the biased surface without top-down symmetry (mov2.mp4 in SI), while it is free when the biased is absent (mov2nobias.mp4 in SI).

Hence, the thermodynamic state point controls how much the adsorped IDP collapses or coils. Furthermore, it is reasonable to suppose that other relevant control parameters are the biomolecule and interface hydrophobicities, although we do not vary them here.

#### Desorption in the coil state at low *T*

The BF model shows that in bulk, at low enough *T* and appropriate *P*, the coil-to-globule transition is reentrant^30^ (Fig. 4) as seen in experiments, at relatively high pressures, in structured proteins,^95–98^ and unstructured polymers.^99^ Furthermore, recent experiments show that the folded domains of fused in sarcoma (FUS), a protein with low-complexity IDRs, undergo cold denaturation, with implications for its mediation of LLPS.^100^

Here, we observe for the IDP under biased confinement the analogous of the cold denaturation at low *T* and a high enough *P* (Fig. 2), e.g., at (*k*_*B*_*T/*4*ϵ, Pv*_0_*/*4*ϵ*) = (0.050, 0.3) (Fig. 4). This low-*T* unfolding is energy-driven due to the contribution of the hydration HBs to the *ℋ*_TOT_ ^30^ and is unaccessible if the energy gain of the HB at the hydrophobic interface is too small (Fig. S1).

We find that, at the reentrant transition, the IDP often extends and desorbs from the biased hydrophobic interface (Fig.5 e, f, and mov3.mp4 in SI). Intriguingly, the IDP flattens out, keeping a characteristic distance from the interface of two layers of water. As a consequence, it minimizes *ℋ*_TOT_ by maximizing the 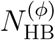 at the two hydrophobic interfaces–the IDP and the wall–and the bulk *N*_HB_.

This observation is consistent with atomistic simulations showing that the bilayer is the most stable free-energy minimum for water confined in a hydrophobic slit pore.^101^ Furthermore, this minimum is energy-driven by the water HBs that saturate to their maximum number per molecule.^86^ Therefore, the BF model captures the atomistic features of the energy-driven double-layer of water while showing the low-*T* flattening of the IDP and its desorption from the interface.

Interestingly, simulations of coarse-grained hydrophobic IDPs in implicit water with effective (water-mediated) *T* -dependent interactions display an upper critical solution temperature (UCST) and a lower critical solution temperature (LCST),^102^ as in experiments with designed IDPs.^103^ Here, the BF model with the reentrant coil-to-globule transition for a hydrophobic IDP offers an ideal test for this phenomenology without introducing effective *T* -dependent interactions, being transferable and water-explicit.

#### Desorption in the coil state under pressurization

At large *P*, e.g., (*k*_*B*_*T/*4*ϵ, Pv*_0_*/*4*ϵ*) = (0.075, 0.5), the Gibbs free energy, Eq.(6), is dominated by the volume term. As discussed for the bulk case^30^ and in agreement with the experiments for protein *P* -induced unfolding,^104^ the large compressibility of the hydration water at hydrophobic interfaces allows the system to reduce the *V*_TOT_ under pressurization. Hence, the protein undergoes a density-driven transition from a globule to a coiled state, as shown by the high-density regions we find around the IDP under these thermodynamic conditions (Fig.5 g, h and Fig. S2 e, f). Furthermore, the high *P* induces a decrease in water HBs number,^60^ diminishing the effective hydrophobic attraction between the surface and the IDP, leading to desorption even in the biased case (mov4.mp4 in SI).

This finding calls for experiments on the protein corona formation and evolution onto nanoparticle and nanomaterials under pressure changes. While *T* effects are known in the corona composition,^105^ to our knowledge, no studies are available as a function of pressure.

#### Adsorption in the coil state under tension

Under tension, e.g., (*k*_*B*_*T/*4*ϵ, Pv*_0_*/*4*ϵ*) = (0.075, −0.2), we find that the IDP unfolds but is still adsorbed onto the biased hydrophobic surface (Fig.5 i, j, and mov5.mp4 in SI). From Eq.(6), we observe that the Gibbs free energy in this thermodynamic regime is minimized by maximizing the volume. From the definition of *V*_TOT_, we note that this condition corresponds to maximizing both *N*_HB_ and 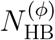 at *P <* 0. Therefore, the IDP loses its globule state to expose the hydrophobic residues to hydration. However, the *P <* 0 unfolding occurs only if the energy gain at the hydrophobic hydration is large enough. Indeed, for the unbiased case with small Δ*J*^(*ϕ*)^*/J* (Table S1) the unfolding at negative *P* is not accessible (Fig. S1).

Under tension, the degree of unfolding is moderate compared to the other cases (at low-*T*, high-*P*, or high-*T*) because a large stretch of the IDP would imply an increase of hydration water with large compressibility, inducing a decrease of *V*_TOT_. Consequently, the protein explores conformations that compromise between globular and unfolded regions.

At the same time, the increase in the number of HBs implies a strengthening of the water-mediated hydrophobic attraction between the IDP and the surface and the consequent adsorption onto the wall. This effect is also evident in the unbiased case. The protein diffuses slowly but, once near the surface, adsorbs irreversibly within our simulation time (mov3nobias.p4 in SI for the protein at (*k*_*B*_*T/*4*ϵ, Pv*_0_*/*4*ϵ*) = (0.15, −0.1) under confinement with top-down symmetry).

Therefore, the unfolding and adsorption of the IDP are enthalpy driven. These observations are particularly relevant in the force-induced protein unfolding and the LLPS under mechanical stress. Cells are permanently exposed to stress resulting from mechanical forces such as, e.g., the tension generated inside adherent and migrating cells, sufficient to unfold cytoskeleton proteins.^106^ Under these tensile conditions, the unfolded proteins can aggregate,^53^ interfering with essential cellular processes and causing severe pathologies–such as neurodegenerative diseases and dementia^107^–for which mechanopharmacology is emerging as a possible control strategy.^108^

## Conclusions

We study a coarse-grained hydrophobic homopolymer chain in a hydrophobic slit pore as a minimal model of an IDP near an interface. We use the BF model in explicit water and perform Monte Carlo free energy calculations under different thermodynamic conditions in confinement with and without top-down symmetry, the latter case mimicking a drift or a weak force pushing the protein toward the interface without limiting its lateral diffusion. Our results reveal that the biased hydrophobic walls drastically affect the coil-to-globule transition of the IDP, reducing its stability region and shifting it to lower *T* and *P*.

We find an intriguing interplay between the surface adsorption-desorption and the coil-to-globule transition of the IDP. A protein unfolds partially when it approaches the surface.^17,21^ However, the IDP can adsorb onto the hydrophobic interface, keeping, at least in part, a globule conformation consistent with recent *in vitro*^24^ and *in vivo* experiments.^94^

At high *T*, the entropy drives the unfolding of the IDP, but not necessarily its desorption when the bias is present. This result is of particular interest in developing strategies based on hyperthermia with protein-functionalized magnetic nanoparticles brought, under the action of forces resulting from external magnetic fields, to high *T* for local treatments of, e.g., cancer cells.^109^

A similar result is also valid when the IDP is under (mechanical) tension. It unfolds but does not necessarily desorbs from the surface. Under these circumstances, the IDP has a less extended conformation, where elongated regions intercalate small globules, keeping their adhesion to the interface. Understanding this mechanism could be crucial to treat diseases involving junctions^110^ as, e.g., cardiac disorders,^111^ where mechanical forces trigger the loss of tertiary and secondary structural elements within anchoring proteins.^106^

Under high-pressure and, possibly, low-temperature stresses, IDPs lose their globular state and desorb from the hydrophobic interface, typically separated by a water bilayer, driven by water’s density at high *P* and water’s energy at low *T*. The energy-driven low-*T* behavior is consistent with atomistic simulations showing that the bilayer is the most stable free-energy minimum for hydrophobically confined water.^101^ Also, it offers an ideal test with a transferable and water-explicit molecular model for recent *in vitro* experiments^103^ and coarse-grained implicit-water simulations, with effective *T* -dependent interactions, of IDPs displaying LLPS with UCST and LCST.^102^ At the same time, our predictions call for new experiments on the protein corona evolution on nanomaterials under pressurization.

## Supporting Information

Additional results for the biased confinement (MP4 movies mov1 - mov5), and the unbiased confinement (MP4 movies mov1nobias - mov3nobias; Table S1; Figures S1 - S3).

## Acknowledgement

We want to acknowledge Prof. Pablo G. Debenedetti for the fruitful discussions about water and biological systems over the years. G.F. acknowledges support from the Spanish grants PGC2018-099277-B-C22 and PID2021-124297NB-C31, funded by MCIN/AEI/ 10.13039/ 501100011033 and “ERDF A way of making Europe”, and the Visitor Program of the Max Planck Institute for The Physics of Complex Systems for supporting a six-month visit started on November 2022.

## Supporting Information

**Table S1:**
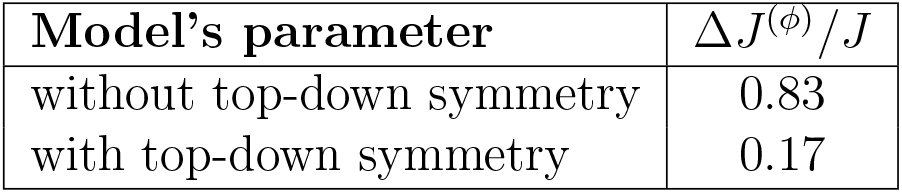
Model’s parameters with or without top-down symmetry. Those not indicated in the table are the same in both cases and are: *v*_HB_*/v*_0_ = 0.5, *J/*4*ϵ* = 0.3, *J*_*σ*_*/*4*ϵ* = 0.05, *k*_1_ = *v*_0_*/*4*ϵ*, with units 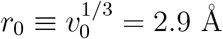 ϵ = 5.8 kJ/mol.

**Figure S1:**
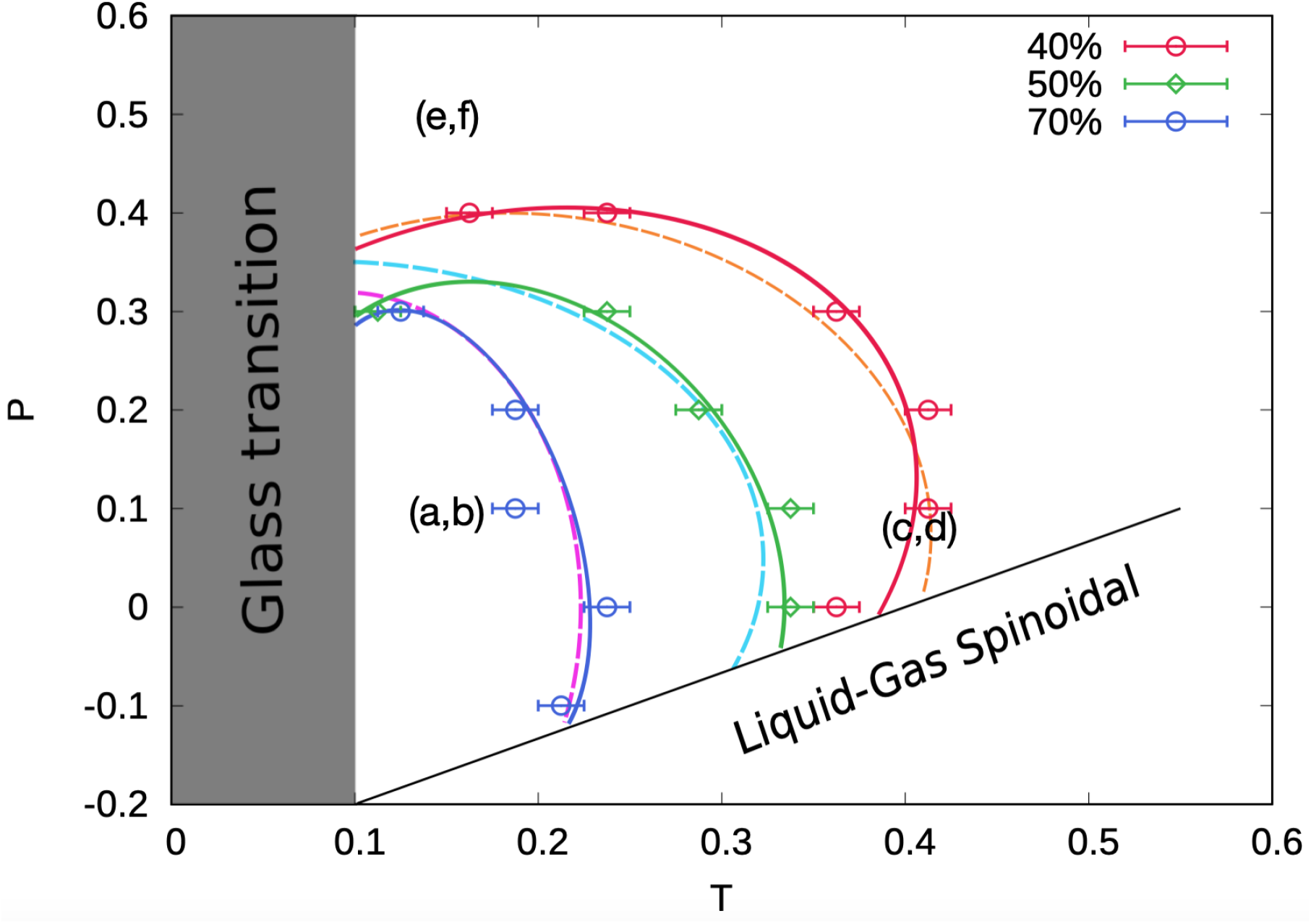
As in Fig.4 but without the bias. Lines are guides for the eyes. The change in liquid-gas spinodal is the same with or without the bias. On the other hand, the change in protein SR without bias is negligible. Lines and symbols are as in Fig.4. Because we reduce Δ*J*^(*ϕ*)^*/J* in the case without bias (Table S1), the protein SR is less accessible compared to Fig.4. The labels (a,b), (c,d), etc., refer to the state points discussed in Fig. S2.

**Figure S2:**
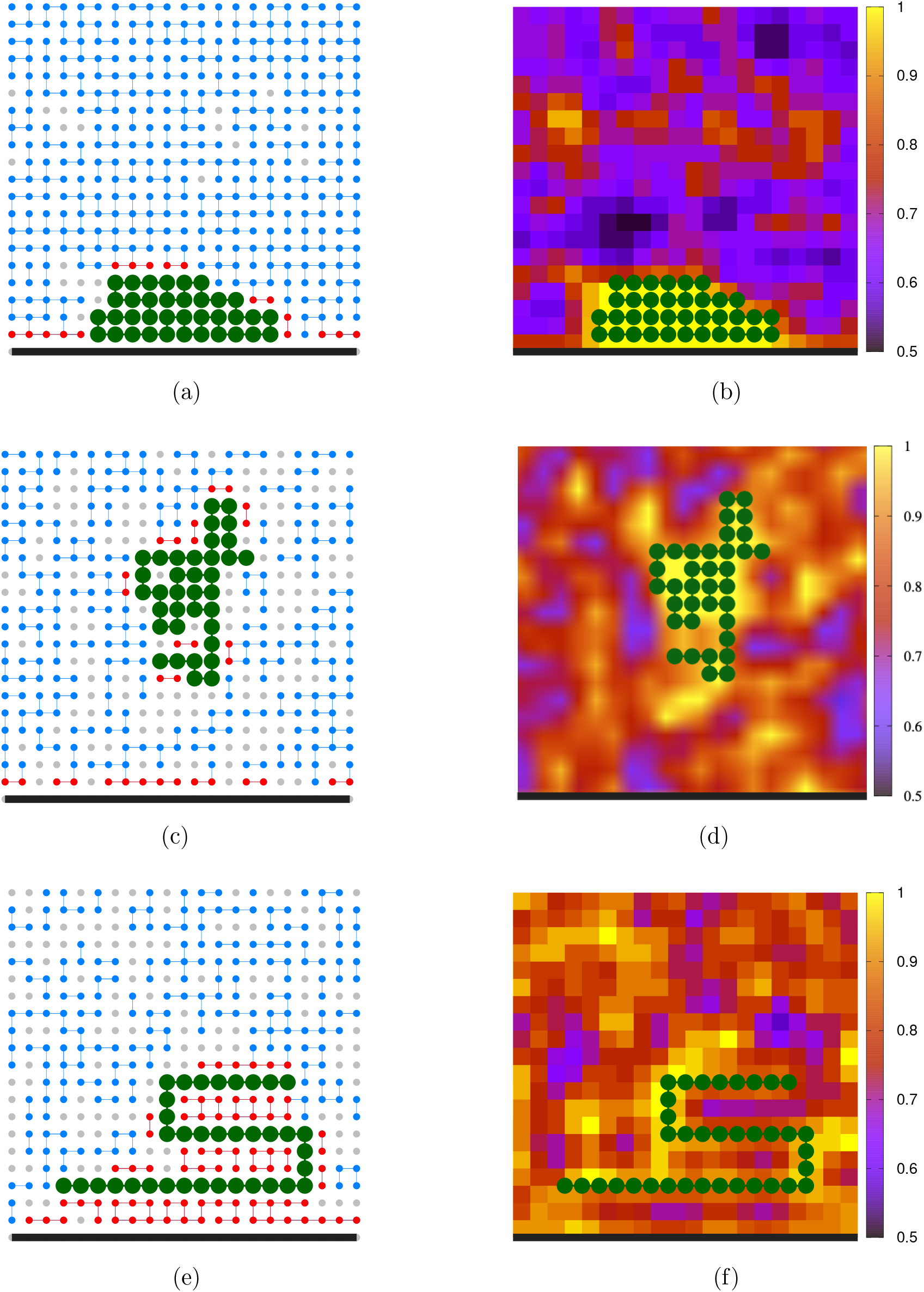
As in Fig. 5 but without the bias. The thermodynamic state points (*k*_*B*_*T/*4*ϵ, Pv*_0_*/*4*ϵ*) of the panels are reported in the phase diagram in Fig. S1: **(a), (b)** (0.15, 0.10); **(c), (d)** (0.40, 0.10); **(e), (f)** (0.15, 0.50).

**Figure S3:**
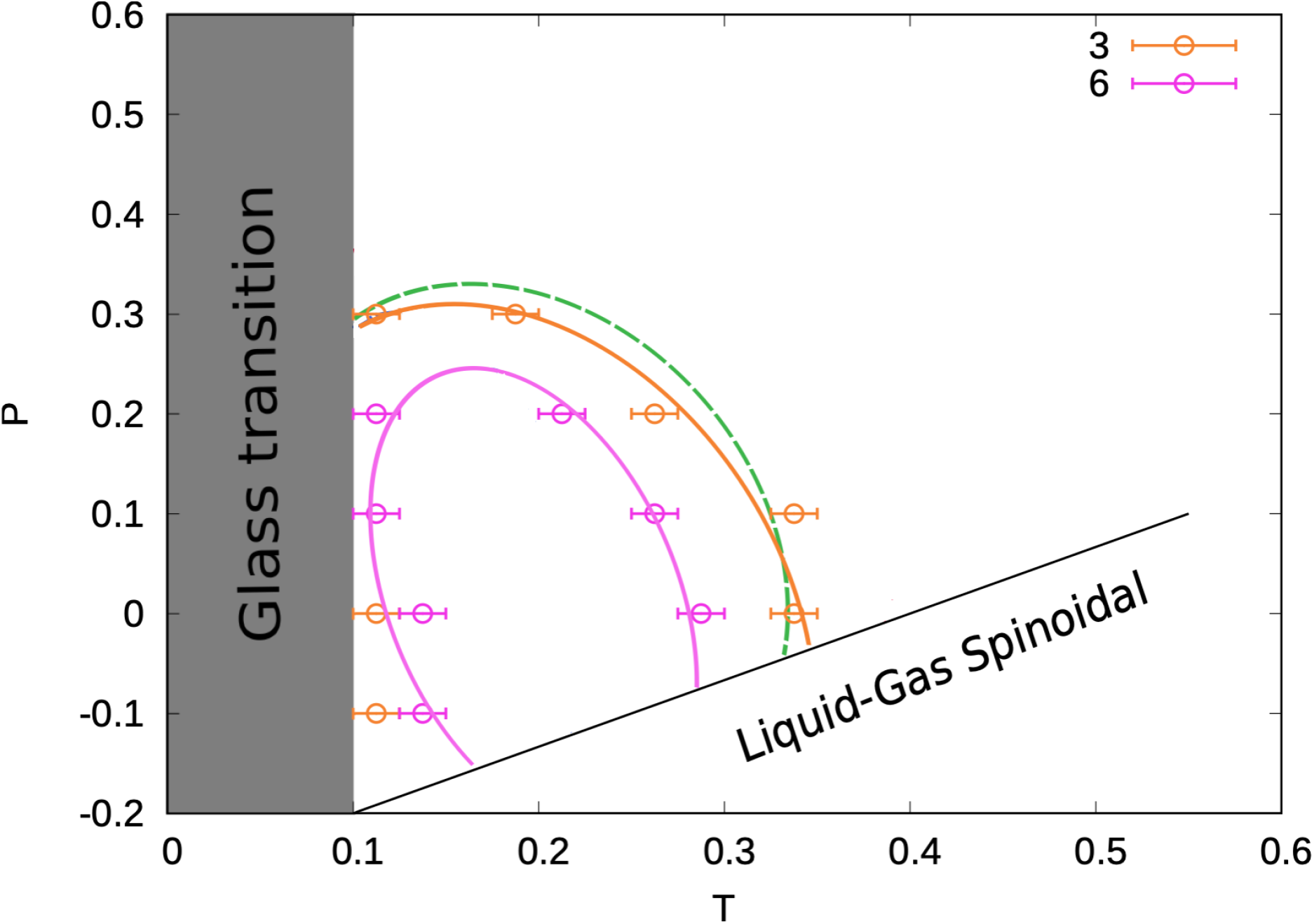
Regions where the protein-interface average number of contact points is *>* 3 (orange symbols and line) and *>* 6 (indigo symbols and line) compared to the protein SR at 50% collapse (dashed green line from Fig. S1) for the confined case without bias. Lines are guides for the eyes. The protein adsorbs when it collapses in the unbiased case. However, at *T* near the glassy state and *P* ≤ 0, the extremely slow protein diffusion can prevent it from reaching the interface and adsorbing onto it.

